# Going beyond rote auditory learning: Neural patterns of generalized auditory learning

**DOI:** 10.1101/2020.12.30.424903

**Authors:** Shannon L.M. Heald, Stephen C. Van Hedger, John Veillette, Katherine Reis, Joel S. Snyder, Howard C. Nusbaum

## Abstract

The ability to generalize rapidly across specific experiences is vital for robust recognition of new patterns, especially in speech perception considering acoustic-phonetic pattern variability. Behavioral research has demonstrated that listeners are rapidly able to generalize their experience with a talker’s speech and quickly improve understanding of a difficult-to-understand talker without prolonged practice, e.g., even after a single training session. Here, we examine the differences in neural responses to generalized versus rote learning in auditory cortical processing by training listeners to understand a novel synthetic talker using a Pretest-Posttest design with electroencephalography (EEG). Participants were trained using either (1) a large inventory of words where no words repeated across the experiment (generalized learning) or (2) a small inventory of words where words repeated (rote learning). Analysis of long-latency auditory evoked potentials at Pretest and Posttest revealed that while rote and generalized learning both produce rapid changes in auditory processing, the nature of these changes differed. In the context of adapting to a talker, generalized learning is marked by an amplitude reduction in the N1-P2 complex and by the presence of a late-negative (LN) wave in the auditory evoked potential following training. Rote learning, however, is marked only by temporally later source configuration changes. The early N1-P2 change, found only for generalized learning, suggests that generalized learning relies on the attentional system to reorganize the way acoustic features are selectively processed. This change in relatively early sensory processing (i.e. during the first 250ms) is consistent with an active processing account of speech perception, which proposes that the ability to rapidly adjust to the specific vocal characteristics of a new talker (for which rote learning is rare) relies on attentional mechanisms to adaptively tune early auditory processing sensitivity.

**Statement of Significance:** Previous research on perceptual learning has typically examined neural responses during rote learning: training and testing is carried out with the same stimuli. As a result, it is not clear that findings from these studies can explain learning that generalizes to novel patterns, which is critical in speech perception. Are neural responses to generalized learning in auditory processing different from neural responses to rote learning? Results indicate rote learning of a particular talker’s speech involves brain regions focused on the memory encoding and retrieving of specific learned patterns, whereas generalized learning involves brain regions involved in reorganizing attention during early sensory processing. In learning speech from a novel talker, only generalized learning is marked by changes in the N1-P2 complex (reflective of secondary auditory cortical processing). The results are consistent with the view that robust speech perception relies on the fast adjustment of attention mechanisms to adaptively tune auditory sensitivity to cope with acoustic variability.

---

A fundamental problem faced by all theories of speech perception is to explain how listeners understand speech despite extensive variability and noise in acoustic patterns across talkers and contexts. One explanation is that listeners overcome acoustic-linguistic variability by tuning the relationship of acoustic cues to linguistic categories by generalizing from talker-specific experiences (for review, see Heald and Nusbaum, 2014; Weatherholtz and Jaeger, 2016). However, many studies investigating the neural correlates of generalized learning in such settings have focused on generalization acquired after long-term rote training (Ross and Tremblay, 2009; Tremblay et al., 2014). While rote training may be one way to rapidly learn the meaning associated with a small set of acoustic patterns (Fenn et al., 2013), it is not the most effective way of producing generalization (Fenn et al., 2013; Greenspan et al., 1988). Instead, broad exposure to a variety of patterns quickly promotes rapid learning of general perceptual categories particularly for speech (cf. Heald and Nusbaum, 2014)

The ability for listeners to generalize beyond their perceptual experiences to novel but related acoustic patterns has been shown to depend on the type of experience or training a learner is given. When participants were trained on a difficult-to-understand computer-generated (synthetic) talker, listeners generalized beyond the words in the training set depending on whether participants were given all novel words during training (generalized training) or if they were given a highly repeated small set of words that repeated (rote training) (Greenspan et al., 1988). Participants who never heard the same word twice during training demonstrated significantly better generalization to untrained words compared to participants who were trained on the small set of repeated words. These results demonstrated that equal amounts of rote and generalized learning lead to different outcomes and thus may be mediated by different processes: The nature and organization of the training materials modulates the degree to which rote or generalized learning takes place. Furthermore, Fenn et al. (2013) found that generalized learning shows clear benefits of memory consolidation during sleep whereas rote learning does not show this benefit, suggesting that there are different neural mechanisms at work in generalization and rote learning.

Thus, if there are different neural systems underlying rote and generalized learning, research using rote learning may not explain how we are able to quickly understand speech from unfamiliar talkers. This conjecture of different neural processes underlying rote and generalized learning has not been explicitly tested.

The evidence that type of training (rote vs. generalized) can determine the degree to which learning will transfer beyond previous experiences raises questions as to what cognitive and neural mechanisms allow for the rapid transfer of learning or generalization. While the neural underpinnings of rapid generalized learning have, to our knowledge, not been empirically examined, rapid generalized learning has been described cognitively as being dependent on the mechanism of selective attention. From a cognitive view, rapid generalized learning of a talker selectively orients attention towards phonetically (classification) relevant acoustic cues and away from irrelevant ones (Francis and Nusbaum, 2002; Goldstone, 1998; Nosofsky, 1986). The emphasis on selective attention in the context of learning a difficult-to-understand talker, as opposed to one that emphasizes say, learning new perceptual *categories*, stems from the idea that adult listeners already possess a complete phonological category system (Chomsky and Halle, 1968; Liberman, 1970). As such, a listener adjusting to a new talker via generalized learning – provided they are speaking the same language – may involve a narrowing of attention towards the most diagnostic acoustic-phonetic cues for the given talker and away from uninformative ones. In these studies, participants are asked to learn to recognize the speech of a difficult-to-understand synthetic talker, which can be thought of as similar to learning to understand a highly idiosyncratic or accented talker, where prior to training, recognition is effortful and difficult (Francis and Nusbaum, 2009). This is because while difficult-to-understand synthetic speech represents a complete orthographic to phonetic system complete with coarticulation, its cue structure is impoverished and contains misleading acoustic cues (Nusbaum and Pisoni, 1985; Schwab et al., 1985). As such, difficult-to-understand synthetic speech possesses greater acoustic-phonetic ambiguity than found in natural speech. This means that prior to learning, there will be more possible phonetic interpretations for a given acoustic pattern than in natural speech. Following rapid generalized learning however, a selective attention account suggests that listening will be much less effortful for new words said by the same talker, as listeners are able to shift attention to the subset of acoustic features that are most diagnostic of the phonemes produced by the talker (Francis et al., 2000). For this reason, training on synthetic speech offers a unique way to investigate how selective attention works in the context of perceptual learning.

Neuroimaging studies investigating the neural correlates of selective attention have shown that instruction to selectively attend to phonetic content modulates activity in anterior parts of the auditory cortex while instruction to selectively attend to a spatial location modulates posterior activity (Ahveninen et al., 2006; Petkov et al., 2004; Woods et al., 2009). This dissociation – likely related to the “what” and “where” processing streams proposed by Raushecker & Tian (2000) – is mirrored in electrophysiological studies demonstrating that the combined activity of these two sources (anterior and posterior) contributes to the morphology of the N1, a negative peak around 100ms in the auditory evoked potential (McEvoy et al., 1997). The electrophysiological measures indicate that activity in anterior parts of the auditory cortex has a longer latency than then activity arising from the posterior source, which has been argued to reflect that the process of identifying an object takes longer than recognizing its spatial origin (Picton, 2011, p. 379). As such, while N1 as a whole has been argued as a marker of attention, the more posterior, earlier-latency N1 source appears to support the gating of awareness to novel sounds, while the more anterior, later-latency N1 source supports a subsequent attentional focus to acoustic features comprising the auditory object (Gutschalk et al., 2008b; Jääskeläinen et al., 2004; Tiitinen et al., 1994). This differentiation between early and late N1 sources relates to earlier work by McCallum and Curry (1979) who argued that the N1 wave should be differentiated into separate waves. While McCallum and Curry (1979) proposed that the N1 wave should be approached as three separate waves, an N1a wave (with a frontotemporal maximal peak at ~70ms), an N1b wave (with a vertex maximal peak at ~100ms, and N1c wave (with a temporal maximal peak at ~140ms), Picton (2011, pp. 379–380) has speculated that the N1c wave can be recognized as arising from this anterior, longer-latency N1 source. Given that in the context of rapid generalized perceptual learning selective attention is thought to exclusively alter how attention is aligned to featural information in phoneme recognition, we hypothesize that the rapid generalized learning of a synthetic talker may exclusively alter the longer-latency N1 activity.

While neural studies on selective attention offer context to our current question, research on the neural correlates of auditory perceptual learning can also offer additional insight. However, it is important to note that generalized perceptual learning of synthetic speech marks a departure from other paradigms used to study the neural underpinnings of perceptual learning. First, extant neural studies investigating perceptual learning have almost exclusively focused on rote (not generalized) rapid perceptual learning, in which participants are repeatedly trained and tested on the same, small set of stimuli. Second, extant perceptual learning paradigms participants have been required to either learn (1) to differentially label sounds that are functionally equivalent in their native language (e.g. Ross and Tremblay, 2009; Tremblay et al., 2014) or (2) to separately label two concurrently presented vowels (with the same spatial origin) (e.g. Alain et al., 2007; Alain and Snyder, 2008; Reinke et al., 2003). Neither of these paradigms examine learning that occurs at the phonological system level as one tries to understand a novel talker but rather focus on the learning of tokens -- either labeling nonnative tokens (new phonological categories) or labeling two stimuli at once--distributing attention over known tokens in novel ways. For this reason, past paradigms that have been used to understand perceptual learning may only offer partial clues into what neural mechanisms support rapid generalized perceptual learning.

Perceptual learning paradigms where participants are given experience with a novel phonetic contrast not in their native language, have documented that learning is marked by an overall decrease in N1 amplitude (maximal at vertex ~100ms) (Alain et al., 2010; Ross and Tremblay, 2009). Work in this paradigm however has argued that this N1 change in this context may be a consequence of habituation and not learning, as the stimulus set only consists of two sounds played repeatedly, often as participants passively listen (Tremblay et al., 2014). However, in more active tasks, in which participants are asked to rapidly learn to segregate concurrently presented vowels, learning has been demonstrated to lead to decreased N1c amplitude (Alain et al., 2007; Alain and Snyder, 2008). As previously mentioned, Picton (2011, pp. 379–380) has argued that an N1c change can be taken as a change in the longer-latency N1 source that supports a subsequent attentional focus to acoustic features comprising the auditory object (Gutschalk et al., 2008a; Jääskeläinen et al., 2004; Tiitinen et al., 1994). As such, this finding suggests that a similar N1c or longer-latency N1 change may be observed following the rapid generalized learning for a difficult-to-understand synthetic talker.

In contrast to rapid generalized learning, rote learning of speech tokens may be more similar to paired-associate learning. Successful paired associate learning entails the formation of associations between stimuli and associated responses rather than the systematic relationships among the speech tokens as a phonological system for a single talker. If rapid rote learning of specific utterances from a difficult-to-understand synthetic talker are characterized by the encoding and the retrieval of episodic memories, it is possible that such learning will not be marked by a change in sensory processing since there is no need to develop a systematic relationship between the talker’s phonetic idiosyncrasies and the native phonological system. This only requires “memorizing” the acoustic patterns and responses. Thus, rapid rote learning should not reorganize attention to acoustic-phonetic properties (as long as the utterances can be discriminated) and therefore should not influence longer-latency N1 activity.

Multi-day rote perceptual learning experiments have also reported an increase in the auditory evoked P2 response after two or three days of training (Bosnyak et al., 2004). The change in the auditory evoked P2 response, which occurs around 200 milliseconds post-stimulus (Näätänen and Winkler, 1999; Ross et al., 2013) is thought to reflect a relatively slow learning process, perhaps relating to the consolidation of a featural representation for the non-native phonetic contrast that participants are learning in long-term memory (Ross and Tremblay, 2009; Tremblay et al., 2014). In the context of learning a difficult-to-understand synthetic talker via generalized learning, it is unclear if a change in the auditory evoked P2 response would be observed. If the change in the auditory evoked P2 response found in the multi-day rote perceptual learning experiments marks the consolidation of a newly learned featural representation in long-term memory for the trained contrast, then an auditory evoked P2 change should not be observed following generalized learning of a difficult-to-understand talker if such learning is best explained by an attention reorientation process. However, if the P2 component is sensitive to the formation of an additional featural representation in long-term memory, it suggests that the auditory P2 evoked response may be sensitive to the number of active featural representations serving current recognition, as the evoked P2 response has been shown to increase after new perceptual categories have been formed (Tremblay et al., 2014, 2009). This is consistent with research showing that the P2 response is sensitive to spectral complexity but only for experts, who presumably rely on more featural representations as spectral complexity increases (Shahin et al., 2005) If the auditory evoked P2 response is sensitive to the number of active feature representations that underlie perceptual recognition, then rapid generalized learning of a difficult-to-understand talker may indeed yield immediate effects on the auditory evoked P2 response. Specifically, if rapid generalized learning leads to a reduction in the ambiguity of how acoustic patterns match to linguistic categories by reducing the number of active feature representations required for ongoing perception, we should see an immediate reduction in the auditory evoked P2 response following training. In contrast to generalized learning, it is unlikely that any change would be observed for rote learning of a difficult-to-understand talker since rote learning of a difficult-to-understand talker may rely more on the encoding and the retrieval of episodic memories. Consequently, rote learning of a difficult-to-understand talker should not alter the number of active featural representations nor, by our logic, a change in the auditory P2.

While previous research examining cortical and subcortical evoked activity associated with perceptual learning has focused on changes in the auditory N1 and P2 waves (sometimes referred to collectively as the N1-P2 complex), a late negativity in the auditory evoked potential starting 600ms post-stimulus onset has also been found to be coincident with improved perception following training (e.g. Tremblay et al., 2014). Given that negative deflections in event related potentials are often affiliated with processes related to error correction (e.g. N400, Late Difference Negativity (LDN), and Error Related Negativity (ERN)), the observed late negativity wave post-training may represent the expansion of an error monitoring mechanism that supports learning. This is consistent with perceptual learning models that specify that the reorganization or formation of perceptual categories (which are implicit in nature) should be dependent on a trial-by-trial prediction-error correction process (Ashby et al., 1998; Ashby and O’Brien, 2005). Under this view, increased late negativity in the auditory evoked potential post-training should only be found under circumstances where training leads to changes in the reorganization or formation of implicit categories (such as those that presumably guide perception). For this reason, we hold that post-training related late negativity in the auditory evoked potential should only be found following generalized learning and not rote learning of a difficult-to-understand synthetic talker.

In the present study, we examine how neural responses produced by generalized learning or rote learning of synthetic speech differ. Are the differences in generalized and rote learning of difficult-to-understand speech reflected in patterns of neural activity during early sensory auditory processing (i.e. during the first 250ms) or are they only manifest in temporally later mechanisms? Specifically, we used a Pretest-Training-Posttest design while performing electroencephalography (EEG). We trained participants using either (1) a large inventory of words in which no words repeated across the experiment (generalized learning) or (2) a small inventory of words where words repeated (rote learning). While participants in the rote learning condition can adopt a simple memorization strategy, subjects in the generalized learning condition cannot use such a strategy as no words repeat across the experiment. Using 128 electrodes and nonparametric significance testing using a permutation test procedure, we compared auditory evoked potentials from stimuli at Pretest to those at Posttest for each learning condition. While our primary hypotheses focused on changes within the auditory N1 and P2 responses (up to 300ms), we assessed differences up until 800ms post word onset, allowing us to determine whether there were any later changes in the auditory evoked potential due to each type of learning.

## 2. Method

### 2.1 Participants

Twenty-nine individuals participated in the generalized learning portion of the experiment (M = 21.4 years, SD = 3.94 years, age range: 18-38, 13 female, 2 left-handed) and thirty-three participants participated in the rote learning portion of the experiment (M = 20.3 years, SD = 2.19 years, age range: 18-26, 17 female, 5 left-handed). All participants were recruited from the University of Chicago and surrounding community. Participants were paid or granted course credit for their participation. All participants identified as native English speakers, with no reported history of either a hearing or speech disorder. Upon completion of the task, all participants reported no prior experience with the stimuli heard in the experiment. Additionally, informed consent was obtained from all participants and the research protocol was approved by the University of Chicago Institutional Review Board.

### 2.2 Stimuli

Stimuli consisted of five hundred monosyllabic words produced using the text-to-speech synthesizer, Rsynth (Ing-Simmons, 1994). These words were taken from phonetically balanced lists that approximate the distribution of phonemes in American English (Egan, 1948) and included nouns, adjectives, verbs, and adverbs. For the generalized learning condition, two test lists (100 words each: Test1 and Test2), and a training list (300 words) were constructed from the synthesized set of five hundred words. For the rote learning condition, 20 words were picked from the synthesized set of five hundred words to serve as the test (both Pretest and Posttest) and Training words. The same 20 words were used for each participant in this condition. The two tests for generalized learning were piloted to be performance balanced in terms of difficulty. While no words repeated across test and training in the generalized learning condition, there was repetition of words in the rote learning condition such that the 20 selected words repeated 5 times each during testing (100 items per test) and 15 times each during training (300 training items). Rsynth uses a formant synthesizer (Klatt, 1980) together with relatively primitive orthography-to-speech rules and it has a reduced and degraded acoustic-phonetic cue set with low acoustic cue covariation compared to natural speech. For these reasons, the intelligibility for Rsynth is quite low. However, listeners show rapid improvement even after a one-hour training session, even when no words repeat across the experiment, increasing their understanding on average by 15 words (15 % improvement) compared to their performance at Pretest (Fenn et al., 2003, 2013; Schwab et al., 1985).

### 2.3 Procedure

Before beginning the experiment, informed consent was obtained. Over the course of the experiment, stimuli were presented binaurally using MATLAB 2015 with Psychtoolbox 3 over insert earphones (3M E-A-Rtone Gold) at 65-70 dB SPL. All participants were initially tested (Pretest), trained (Training), and retested (Posttest) on their identification performance for monosyllabic synthetic speech stimuli. In the generalized learning condition, the test lists used at Pretest and Posttest were counterbalanced across participants to ensure that any change in performance from Pretest to Posttest would reflect learning. As the testing material was the same at Pretest (5 repetitions for each word--100 test trials), Training (15 repetitions for each word--300 training trials) and Posttest (5 repetitions for each word--100 test trials) for the rote learning condition, there was no need to counterbalance the tests for the rote learning condition. For both rote and generalized learning, test trials consisted of participants hearing a synthetic speech token and, after a short delay, being asked to type back what they heard. For training, this identification procedure was followed by visual written feedback in tandem with an additional auditory presentation of the synthetic speech token. In the generalized learning condition, no words ever repeated across the experiment (i.e., participants heard 100 unique words during the Pretest, 300 unique words during Training, and 100 unique words during the Posttest). In the rote learning condition, the same 20 selected words were repeated five times each during both the Pretest and Posttest and 15 times each during Training (randomized in blocks of 20). At the conclusion of the experiment, participants in the rote condition were given a generalized learning test of 100 novel words (identical to Test 1 in the generalized learning condition). This was done to replicate previous research that showed that rote learning training leads to poorer generalization to untrained words compared to training on all novel words when learning a difficult-to-understand talker (Fenn et al., 2013; Greenspan et al., 1988). Important to our present hypotheses here, these previous studies show that rote learning training for 20 repeat words of a difficult-to-understand talker does allow for some generalization to untrained words, but that this generalization is significantly weaker than generalization found for those who are trained on all novel words (Fenn et al., 2013; Greenspan et al., 1988). For this reason, we expect generalization performance for rote individuals to be better than generalization performance found in the generalized learning condition at Pretest, but to be significantly worse than generalization performance found in the generalized learning condition at Posttest. Electroencephalograph (EEG) signals were recorded continuously during the whole experiment. After the experiment concluded, participants’ heads were photographed using a geodesic dome with 11-mounted infrared cameras to precisely determine the location of all 128 electrodes (Russell et al., 2005).

### 2.4 Data Acquisition

Neurophysiological responses were obtained using an Electrical Geodesics, Inc. (EGI) GES 300 Amp system (output resistance 200MΩ with a recording range from 0.01 to 1000Hz). The high-density EEG (128 electrodes) was recorded at 250 samples/second in reference to vertex using unshielded HCGSN 130 nets. Before both Pretest and Posttest periods, impedances were minimized by reseating or, if necessary, by rewetting electrode sponges using a transfer pipette and saline (to 50 kΩ or less). Resulting amplified EEG signals were recorded using EGI Net Station software (v. 4.5.7) on a computer running Mac OSX (10.6) operating system. No filtering was applied to the EEG signal at acquisition. Trial types were tagged in Netstation using the Netstation Toolbox in Psychtoolbox. Timing of tags was corrected during preprocessing as a mean tagging latency of 15ms (SD: 2.4ms) was found between the stimulus presentation computer running Mac OSX and Net Station via EGI’s audio timing test kit.

### 2.5 EEG Preprocessing

EEG recordings were preprocessed in Brain Electrical Source Analysis Program (BESA Research 7.0). Electrode coordinates from individuals’ net placement photos were used to assign individual sensor locations for each participant. Recordings were filtered with 0.3-50 Hz bandpass and 60 Hz notch filters to remove electrical noise. Voltage was re-referenced to the average of all electrodes. Based on the trial tags, epochs of interest around the times of stimuli presentation were selected as 200 ms before to 800 ms after the onset of the stimulus. Epochs were then examined for artifacts including eye blinks and movements. Beyond visual inspection, voltage threshold detection was also used (voltage thresholds for eye movements were 150 μV for horizontal movements picked up in the EOG electrodes, and 250 μV for vertical movements). Artifacts were removed from the epochs of interest using ocular source components using the brain electrical source analysis software (BESA Research 7.0) (Berg and Scherg, 1994; Picton et al., 2000). In some cases, artifacts due to large movements or to sweat could not be removed by ICA. In these cases, the contaminated trials were not included in further analysis. Individual channels that were problematic for a majority of trials (amplitude >150 μV indicating excessive noise, <0.01 μV indicating low signal, or changes of >75 μV from one sample to the next) were replaced by interpolation using surrounding channels. Because electrode impedances were only checked before Pretest and Posttest periods, we declined to analyze EEG data from the training block to ensure we only present the highest quality data.

The data collected from three participants from the generalized learning condition (2 male, all right handed) and data collected from three participants from the rote learning condition (2 male, all right handed) were removed from further analysis due to excessive artifact contamination (removal of more than 30 trials, or less than 70 remaining trials in a condition). In the generalized learning condition, the remaining twenty-six participants had an average of 90 trials remaining for the Pretest (*SD*: 8.69, range: 74-100), and an average of 88 trials remaining for the Posttest (*SD* 7.22, range: 75-99). In the rote learning condition, the remaining thirty participants had an average of 89 trials remaining for the Pretest (*SD*: 8.10, range: 71-100), and an average of 91 trials remaining for the Posttest (*SD* 8.43, range: 70-100). For each participant, averaged waveforms for the conditions of interest (e.g. Pretest, Posttest) were created, as were corresponding files for topographic analysis in RAGU (Koenig et al., 2011). The 100ms pre-stimulus period was used to baseline correct the ERP averages, by subtracting the average during the pre-stimulus period from each time point in the waveform. To compute topographic maps, participant-specific three-dimensional electrode locations were used. The averaged ERP data, participant-specific electrode location files, and the behavioral data have been made available on Open Science Framework (https://osf.io/kwcsv/?view_only=ee2b903d3c2c467aa6b954b19c7a2afa).

### 2.6 Statistical Analyses

#### 2.6.1 Global Analyses

To conduct global analyses over the course of the entire epoch using every electrode, we used RAGU (RAndomization Graphical User interface:), an open-source Matlab-based program that performs nonparametric significance testing by generating 5000 simulations in which data from the conditions of interest have been randomly shuffled to bootstrap a control data set (see http://www.thomaskoenig.ch/index.php/software/ragu/download). This set of simulations functions as a null distribution, against which the observed data can be compared, usually using a measure of effect size such as global field power (GFP, or the standard deviation across electrodes at a given time point) or global map dissimilarity (GMD, a measure that captures scalp topography differences between conditions at a given time point). This avoids biases associated with a priori assumptions about which time windows and electrodes should be included in analysis (Koenig et al., 2011; Murray et al., 2008). We used RAGU to perform a GFP analysis and a TANOVA (topographic analysis of variance) that compared strength of scalp field potential and scalp topography respectively, between Pretest and Posttest.

RAGU calculates GFP at every time point in the epoch of interest as follows: 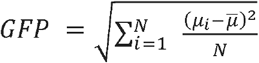, where N is the number of electrodes, *µ_i_* is the voltage of electrode i, and 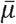 is the mean voltage across all electrodes (Koenig and Gianotti 2009). Thus, GFP is a measure of standard deviation across electrodes. Conceptually, this means if there is a strong response over part of the scalp, the GFP will be greater due to more variance across locations, whereas weak responses will yield low GFPs. GFP also has the benefit of being entirely reference independent. Once the observed GFP is calculated, the data are shuffled between conditions and GFP is recalculated. This reshuffling procedure is carried out 5,000 times at each time point to obtain the null distribution of GFP for a given time point. At each time point, a p-value is calculated that represents the proportion of randomized GFPs that exceed the observed GFP.

While GFP is a good measurement of differences in the strength of potentials across the scalp, potentially important topographical information is lost by calculating standard deviation across all electrodes. RAGU’s TANOVA measures differences in topographical distributions of voltage between pairs of conditions or time points. The measure of effect size used is generalized dissimilarity s across the experimental conditions: 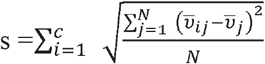, where c is the number of conditions, N is the number of electrodes, 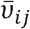 is the mean voltage of condition i at electrode j across subjects, and 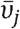 is the mean voltage at electrode j across all subjects, with conditions averaged together (Koenig and Melie-García 2010). Because this measure accounts for differences between condition-wise maps at individual electrodes, it preserves topographical information, unlike GFP: the farther apart the voltage at electrode j in condition A versus B, the larger the squared difference added to s, and therefore the larger the difference in voltage patterns on the scalp.

Once RAGU has calculated the generalized dissimilarity s from the data, it shuffles the data between conditions and re-calculates s 5,000 times to generate a null distribution of generalized dissimilarities. These are the effect sizes that would be expected in the absence of a true difference between the conditions. A p value is calculated at each time point by comparing this null distribution to the observed data as with GFP. The TANOVA was used to identify periods of interest as time windows where there appeared to be significant map differences between Pretest and Posttest (p<0.05). Before the TANOVA analysis, we normalized data by dividing all voltage values of a given map by its time-specific GFP. This was done so that significant differences found between the conditions in the TANOVA analysis could be attributed solely to underlying differences in source contributions in the brain.

For both GFP and TANOVA analyses, we also report whether the window passed a duration threshold test. This was done by collecting the duration of continuous windows found to be significant in the bootstrapped data. The distribution of these durations represents the distribution of duration under the null hypothesis that the data are interchangeable between conditions, as they were obtained from the shuffled data (in which the data are interchanged between conditions randomly). For each test, we set the threshold for the duration as the 95th percentile of spurious window durations that appear across the 5,000 random permutations. Windows in the observed data that pass this threshold testing are clearly noted; however, we decided to report all windows, especially those before 300ms, given the transient nature of the N1 and P2 auditory evoked potentials (Picton, 2011) of interest to us here.

Beyond the RAGU analysis, we used BESA Statistics 2.0 to ascertain which electrodes were responsible for the observed topographic changes. To do this, we averaged each electrode’s voltage over the windows identified in the TANOVA analysis and performed paired sample t-tests between Pretest and Posttest. This analysis used a spatio-temporal permutation-based correction to adjust for multiple comparisons. For these analyses we used a cluster alpha level of 0.05 for cluster building, 5000 permutations, and a channel distance of 4cm that resulted in an average of 6.58 neighbors per channel for the generalized learning condition and an average of 7.09 neighbors per channel for the rote learning condition.

#### 2.6.2 Source Level Analysis

To investigate the intracranial sources underlying the topographic window changes identified by the TANOVA analysis in RAGU we used the local auto regressive average (LAURA) model in BESA Research 7.0. LAURA is a distributed source localization method that does not make a priori assumption with respect to the number of discrete sources. Similar to other distributed volume inverse imaging methods, LAURA seeks to find a solution where the distribution of the current over all source points is minimized while optimally trying to explain the observed topography. LAURA however uses a spatial weighting function to account for the fact that source strength should decrease by the inverse of the cubic distance between a putative source and recording electrodes on the scalp. The result of this technique is a spatio-temporal projection of current density in a neuroanatomical space similar to a functional map in fMRI. LAURA modeling was applied to the full ERP epoch (−200 to 800ms) for each participant. Using a cluster-based permutation test in BESA Statistics 2.0, we computed average distributed source images across the time windows identified by the TANOVA analysis in RAGU. These average distributed source images were then contrasted between Pretest and Posttest for each learning condition using a cluster-based permutation test (Contrast: Posttest-Pretest). This analysis allowed us to identify anatomical locations in the brain volume responsible for the topographical differences identified by the TANOVA analysis. For this analysis, cluster values were ascertained by the sum of all t-values within a given cluster. The significance of observed clusters is determined by generating and comparing clusters from 5000 permutations of the data between stimulus conditions.

Statistically, the results reported from this analysis are highly conservative as BESA corrects for multiple comparisons *across all voxels and time points* in order to control the familywise error rate. Due to the conservative nature of this analysis and that the windows we are investigating have been previously identified through the TANOVA analysis in RAGU, all identified clusters from this analysis are reported including null results.

## 3. Results

### 3.1 Behavioral Results

#### 3.1.1 Generalized learning behavioral results

Word recognition performance at Posttest (number of words transcribed correctly) was subtracted from word recognition performance at Pretest (number of words correct) to obtain a subject-specific learning score. Pretest performance averaged 29 words correct out of 100 (*SD*: 8.19, range: 15 to 45). After training, recognition performance on the posttest significantly increased to an average of 42 words correct out of 100 (*SD*: 13.27, range: 20 to 71) [paired sample t-test: *t*(25) = 7.11, *p* < .00001; Cohen’s *d* = 1.402]. This means that individuals significantly recognized more words at Posttest than they did at Pretest, despite no words repeating across the tests (or training) in the generalized learning condition.

#### 3.1.2 Rote learning behavioral results

Similar to generalized learning, word recognition performance at Posttest (number of words transcribed correctly) was subtracted from word recognition performance at Pretest (number of words transcribed correctly) to obtain a subject-specific learning score. Pretest performance averaged 16 words correct out of 100 (*SD*: 7.83, range: 4 to 30). After training, recognition performance on the posttest significantly increased to an average of 95 words correct out of 100 (*SD*: 8.58, range: 68 to 100) [paired sample t-test: *t* (29) = 41.48, *p* <.00001; Cohen’s d = 7.571]. This indicates that training significantly helped individuals to appropriately recognize the words shown at Pretest by the Posttest in the rote learning condition.

Performance on the additional generalized learning test (all novel words) for those in the rote learning condition shows that participants on average correctly identified 35.7 words correctly out of 100 (*SD*: 8.65, range: 13-47 words). While this performance was better than performance demonstrated by individuals in the generalized learning condition at Pretest [independent, equal variance not assumed two sample (Welch’s) t-test: *t*(53.55) = 2.77, *p* = 0.008; Cohen’s *d* = 0.74], it was also significantly worse than performance by those in the generalized learning condition at Posttest [independent, equal variance not assumed two sample (Welch’s) t-test: *t*(41.90) = −2.18, *p* = 0.035; Cohen’s *d* = 0.60]. This finding replicates previous work showing that while rote training (repeat experience on a small subset of words) can yield some generalized learning, that such generalized learning is significantly weaker than generalized learning that results from training on a set of all novel words.

### 3.2 Electrophysiology Results

#### 3.2.1a Generalized learning electrophysiology results

Figure 1, panel A, shows the Grand Average ERPs (across all subjects in the generalized learning condition) elicited during both Pretest and Posttest. For both Pretest and Posttest, N1 and P2 had maximal voltages at central sites (e.g. C3, Cz, C4). At Pretest, N1 peaked at 112ms at Cz, while P2 peaked at 200ms at Cz. At Posttest, N1 peaked at 108ms at Cz, while P2 peaked at 200ms at Cz. Consistent with prior research, inverted polarity for the N1 and P2 waves was found over sites P9 [left temporal] and P10 [right temporal] (See Figure 1, panel B) below the Sylvian fissure, thereby suggesting that the neural generators for both N1 and P2 are in or near primary auditory cortex (Andrews et al., 1990; Liégeois-Chauvel et al., 1994; Scherg et al., 1989; Yvert et al., 2005).

**Figure 1.**
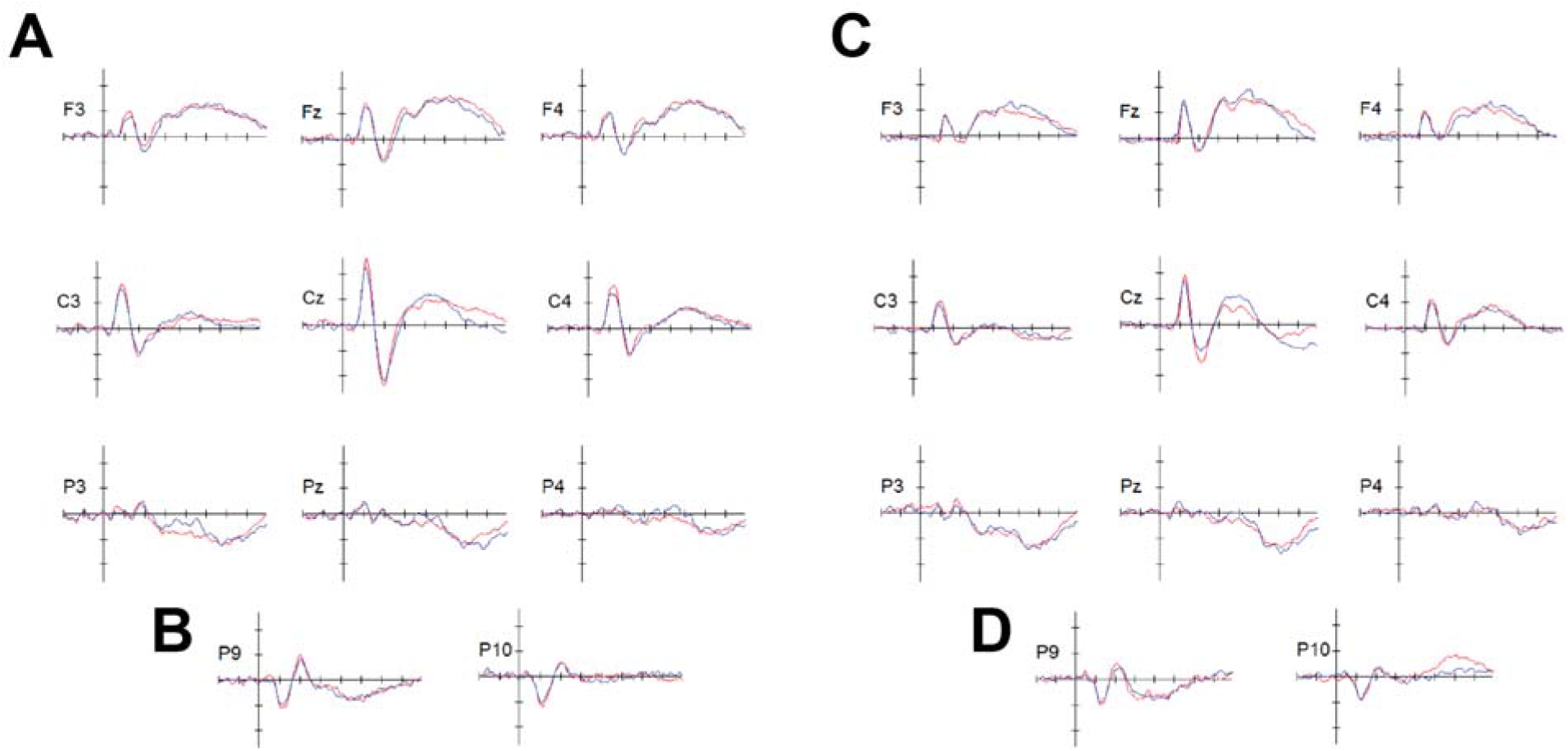
Grand Average ERPs for Generalized Learning (Panel A) and Rote Learning (Panel C) for the nine centralized locations using a virtual montage of the 10-20 system available in BESA Research 7.0 (F3, Fz, F4, C3, Cz, C4, P3, Pz, P4). Grand Average ERPs for P9 and P10 electrodes for Generalized Learning (Panel B) and Rote Learning (Panel D) demonstrate the inversion of the N1-P2 complex that is typical of auditory evoked potentials. The ERPs associated with Pretest trials are shown in red and the ERPs associated with Posttest trials are shown in blue. Horizontal tick marks span 100ms and vertical tick marks represent 1μV, negative is plotting up.

#### 3.2.1b Rote learning *electrophysiology* results

Figure 1, panel C, shows the Grand Average ERPs (obtained by averaging across all subjects in the rote learning condition) elicited during Pretest and Posttest. At both Pretest and Posttest, N1 and P2 had maximal voltages at central sites (e.g. C3, Cz, C4), with the highest peak at Cz. At Pretest, N1 peaked at 112ms at Cz, while P2 peaked at 200ms at Cz. At Posttest, N1 peaked at 108ms at Cz, while P2 peaked at 200ms at Cz. Similar to the generalized learning condition, inverted polarity for the N1 and P2 waves was found over sites P9 [left temporal] and P10 [right temporal] below the Sylvian fissure in the rote learning condition (See Figure 1, panel D) indicating that the neural generators for both N1 and P2 in this condition are also in or near primary auditory cortex (Andrews et al., 1990; Liégeois-Chauvel et al., 1994; Scherg et al., 1989; Yvert et al., 2005).

#### 3.2.2a Analysis of overall amplitude difference for Generalized Learning

Based on analysis with RAGU we identified two windows in the time series of auditory evoked potential in which the observed GFP difference exceeded the top bound of the null distribution’s 95% confidence interval. Both of these windows had sufficient length and passed window thresholding. In the first window from 116ms to 208ms, GFP was shown to be lower at Posttest (mean: 1.16 μV, *SD*: 0.14) compared to Pretest (mean: 1.39 μV, *SD*: 0.14) (See Figure 2, top plot). The mean GFP difference in the observed data in this time window was 0.22 μV (Posttest-Pretest). The mean GFP difference under the null was 0.073 μV, 95% CI [0.03 μV, 0.14 μV]. During this time window, the RAGU GFP procedure showed that on average there was 0.02 probability that the effect size of GFP under the null was larger than the observed difference in GFP (minimum and maximum p-values of where the observed data falls in the null distribution for GFP difference in this interval are respectively, 0.002 and 0.05) (See Figure 2, bottom plot).

**Figure 2.**
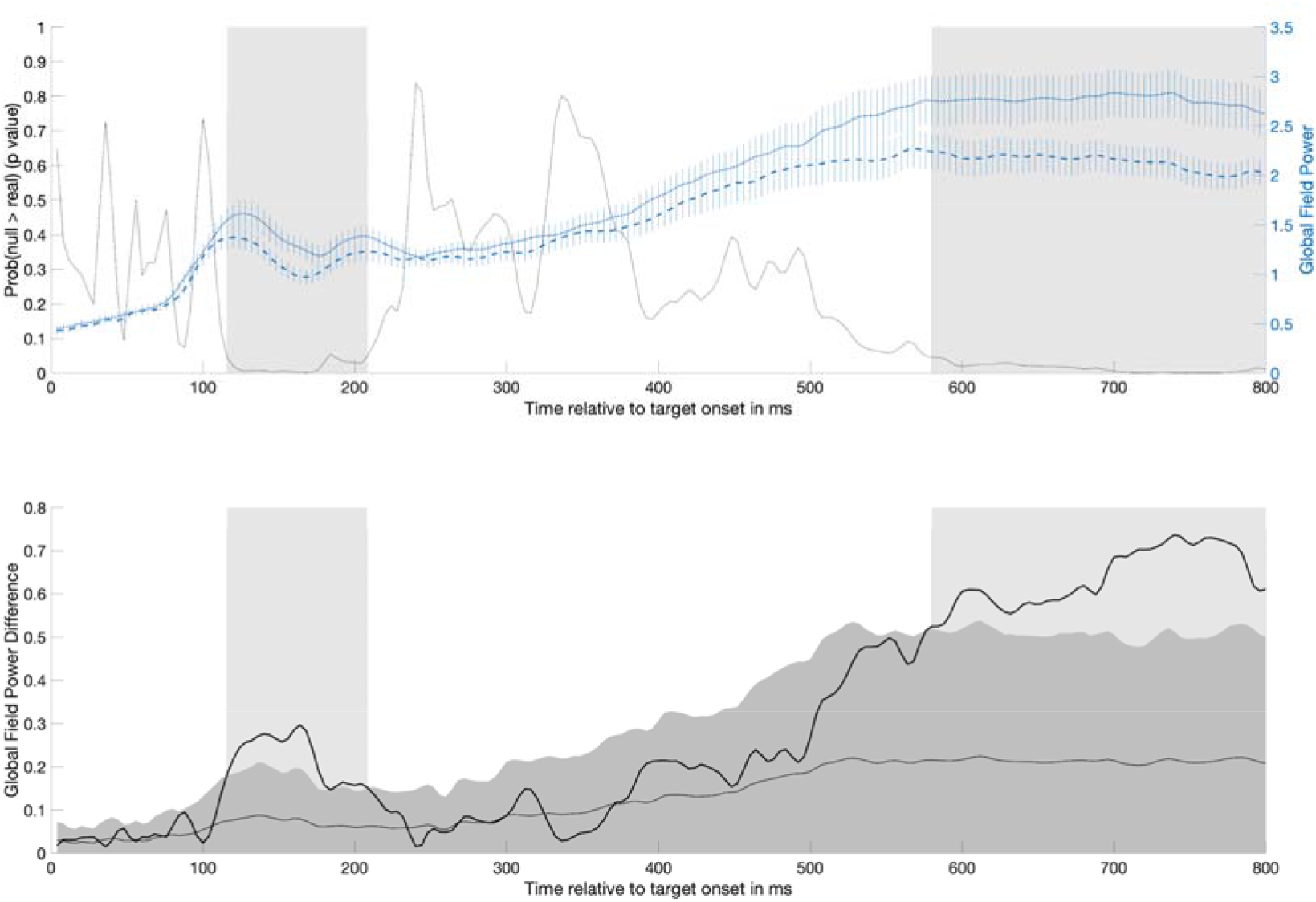
The top plot Mean GFP (right y-axis) over time for both Pretest (short dashed line) and Posttest (long-dashed line) (Error bars show +/- 1SE) in the generalized learning condition overlaid on the probability over time that the GFP difference under the null was larger than the observed difference in GFP (black line) (left y-axis). Significant time periods identified by the GFP analysis are shaded gray: one occurring from 116ms to 208ms and another occurring from 580ms to 800ms. The bottom plot shows the difference in GFP between Pretest and Posttest (dark black line). For context, the mean (light grey line) and 95% confidence interval (dark gray area) for the GFP difference expected due to random chance (estimated from randomizing the data 5000 times) has been plotted. Significant time periods are again shaded gray, although note that these periods are identified by the observed difference in GFP exceeding the upper bound of the 95% confidence interval of the shuffled data.

In the second window from 580ms to 800ms, GFP was shown to be lower at Posttest (mean: 2.14 μV, *SD*: 0.08) compared to Pretest (mean: 2.77 μV, *SD*: 0.05) (See Figure 2, top panel). The mean GFP difference in the observed data in this time window was 0.64 μV (Posttest-Pretest). The mean GFP difference under the null was 0.21 μV, 95% CI [0.05 μV, 0.49 μV]. During this time window, the RAGU GFP procedure showed that on average there was 0.01 probability that the effect size of GFP under the null was larger than the observed difference in GFP (minimum and maximum p-values of where the observed data falls in the null distribution for GFP difference in this interval are respectively, 0.0014 and 0.05) (See Figure 2, bottom panel).

#### 3.2.2b Analysis of overall amplitude difference for Rote Learning

Using the same analysis on the rote learning data yielded no significant GFP differences. The top panel of Figure 3 plots the observed GFP values over time at Pretest and Posttest for rote learning, along with the probability of obtaining a GFP difference by chance more extreme than the observed GFP difference. The only time that the observed data came close to falling in the extreme tail (p < 0.05) of the null distribution is around 300ms. The bottom panel of Figure 3 shows where the observed GFP difference fell relative to the null distribution (mean and 95% confidence interval are shown).

**Figure 3.**
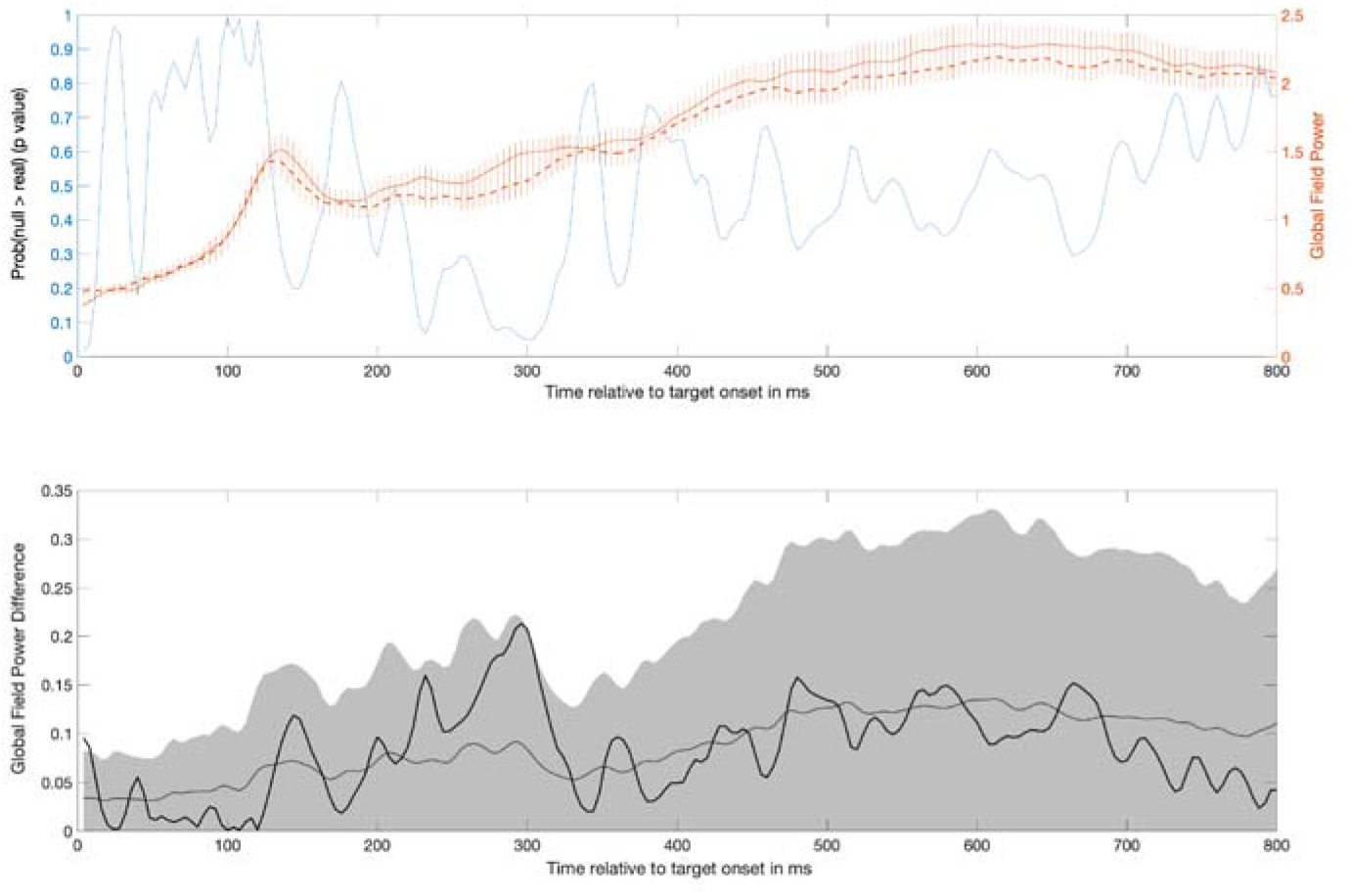
The top plot shows mean GFP (right y-axis) over time for both Pretest (short dashed line) and Posttest (long-dashed line) in the rote learning condition (Error bars show +/- 1SE) overlaid on the probability over time that the GFP difference under the null was larger than the observed difference in GFP (black line) (left y-axis). No significant time windows were observed. The bottom plot shows the difference in global field power between Pretest and Posttest. For context, the mean (light grey line) and 95% confidence interval (dark gray area) for the GFP difference expected due to random chance (estimated from randomizing the data 5000 times) has been plotted.

#### 3.2.3a Analysis of overall topographic difference for Generalized Learning

We used RAGU to generate the Generalized Dissimilarity between Pretest and Posttest that may be obtained due to chance by shuffling the data 5000 times. This distribution for the Generalized Dissimilarity statistic under the null was then compared to the observed Generalized Dissimilarity statistic. For generalized learning, only one window (from 250ms to 272ms) for topographic change was identified (See Figure 3, panel A).

While this interval did not pass the window threshold test in RAGU, it’s appearance at the end of the N1-P2 time window (previous to 300ms) fits our prediction that generalized perceptual learning reorganizes sensory evoked responses. The average observed Generalized Dissimilarity statistic between Pretest and Posttest in this time window was 2.99, while the average Generalized Dissimilarity statistic between Pretest and Posttest under the null was 2.22, with a 95% confidence interval of 1.76 and 2.77. During this time window, the RAGU TANOVA procedure showed that on average there was 0.011 probability that the Generalized Dissimilarity statistic between Pretest and Posttest under the null was larger than the observed difference in the Generalized Dissimilarity statistic (minimum and maximum p-values of where the observed data falls in the null distribution for the Generalized Dissimilarity statistic in this interval are respectively, 0.01 and 0.04) (See Figure 4, panel A).

**Figure 4.**
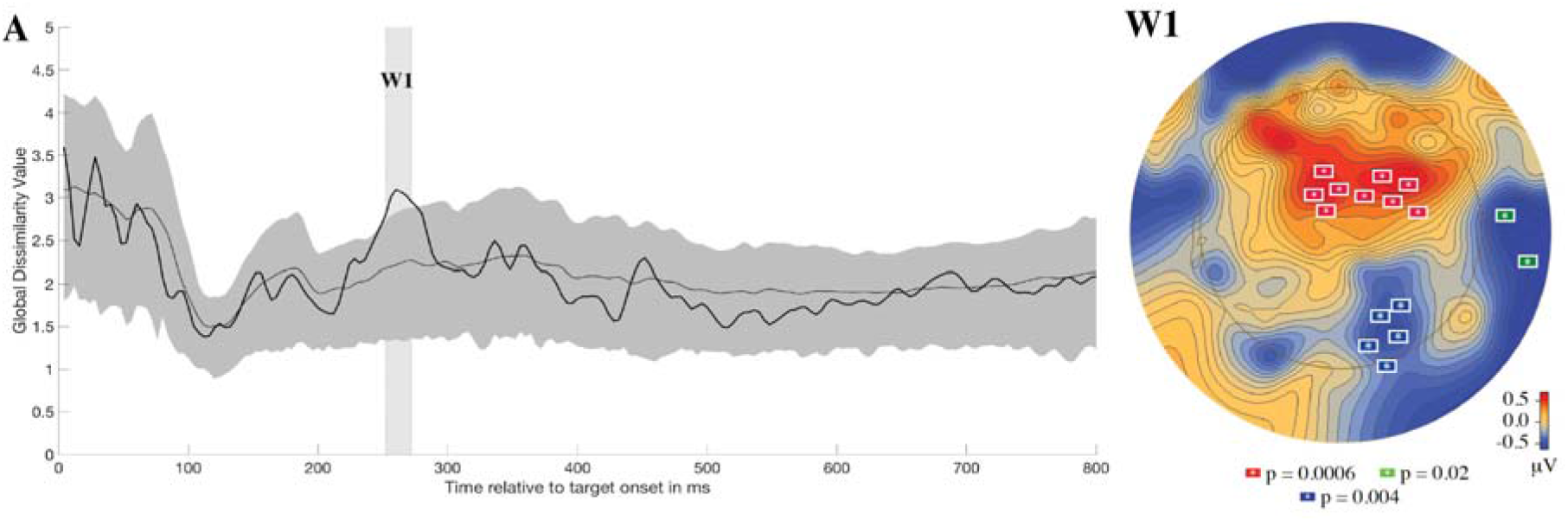
Plot A shows how the Generalized Dissimilarity statistic between Pretest and Posttest topographies varies overtime in the generalized learning condition (black line). For context, the mean (light grey line) and 95% confidence interval (medium gray area) for the Generalized Dissimilarity statistic expected due to random chance (estimated from randomizing the data 5000 times) has been plotted. Plot W1 shows the results of the spatio-temporal permutation-based analysis that was performed on the W1 window found in RAGU. This plot shows the average topographic difference between Pretest and Posttest (Contrast: Posttest-Pretest) and indicates where the three significant electrode clusters (p<0.05) are topographically located.

A spatio-temporal permutation-based analysis performed in BESA Statistics 2.0 on average surface electrode activity during this time window identified three significant electrode clusters driving the topographic change between Pretest and Posttest (See Figure 4, panel W1). The first significant cluster (*p* = 0.0006) was found between frontal and central electrodes left of the midline and comprised the following EGI electrodes (approximate 10-10 equivalent is included if available (Luu and Ferree, 2000)): 6 (Fcz), 7, 12, 13 (FC1), 110, 111(FC4), 112 (FC2), 117(FC6), and 118). The second significant cluster (*p* = 0.0038) was found over the left preauricular region and comprised the following EGI electrodes: 114 (T10) and 113. The third significant cluster (*p* = 0.0154) was found between parietal and occipital electrode left of the midline and comprised the following EGI electrodes (approximate 10-10 equivalent is included if available): 83 (O2), 84, 89, 90 (PO8) and 91.

#### 3.2.3b Analysis of overall topographic difference for Rote Learning

To determine if a change in the scalp distribution of brain electrical activity occurred due to rote learning, we used RAGU to calculate the observed Generalized Dissimilarity statistic between Pretest and Posttest along with this statistic for 5000 shuffles of the data to calculate a null distribution. This analysis identified six windows where the observed Generalized Dissimilarity statistic exceeded the top bound of the null distribution’s 95% confidence interval. However, only two of these windows (window 4 (W4): 424-484ms and window 6 (W6): 660-800ms) passed the window threshold test in RAGU (See Figure 5, Panel A). Table 1 reports (1) the observed Generalized Dissimilarity statistic between Pretest and Posttest, (2) the mean and (3) 95% confidence interval for the Generalized Dissimilarity statistic between Pretest and Posttest under the null, as well as (4) the probability of the observed data given the shuffled data for each of these identified time windows.

**Figure 5.**
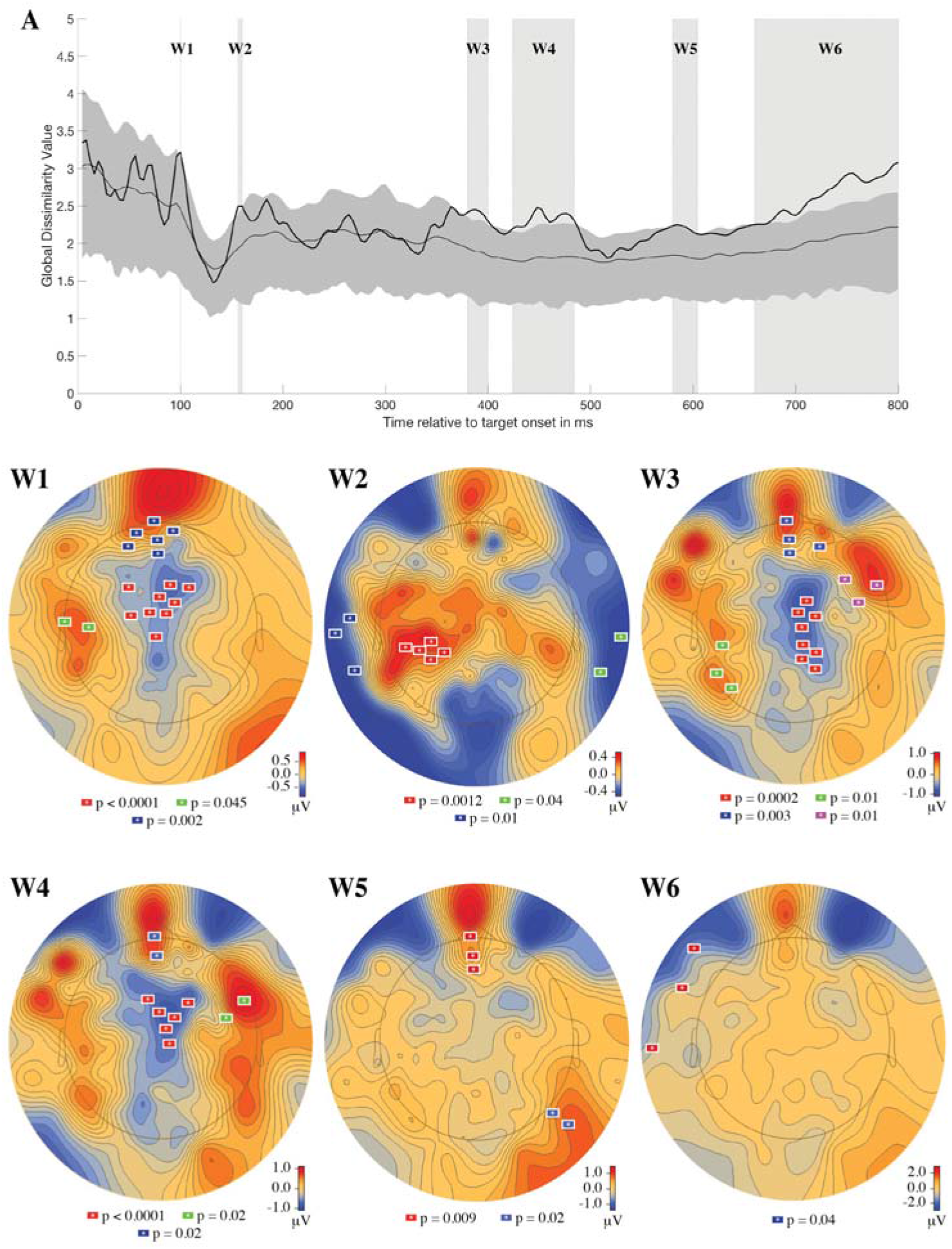
Plot A shows the Time-varying generalized dissimilarity between Pretest and Posttest topographies for Rote Learning (black line). For context, the mean (light grey line) and 95% CI (medium gray area) for the generalized dissimilarity expected due to random chance (estimated from randomizing the data 5000 times) has been plotted. Six windows were identified where the observed data exceeded the upper bound of the 95% CI of the shuffled data. Plots W1, W2, W3, W4, W5, and W6 show the results of the spatio-temporal permutation-based analysis that was performed on each window. All electrode clusters shown, uniquely colored on each plot, are significant at a p<0.05 level.

**Table 1.** Significant time windows showing topographic change from Pretest to Posttest identified by the TANOVA RAGU analysis in the Rote Learning Condition. For each window, the observed generalized dissimilarity (GD) between Pretest and Posttest is reported along with the Mean and the 95% confidence interval (CI) from the permutation distribution. The final column (Probability of the Observed GD statistic (or greater) under the permutation distribution) reports the percent of the 5000 shuffled versions of the data that obtained a GD statistic between Pretest and Posttest more extreme than actually observed. Shaded rows indicate windows that pass the duration threshold in RAGU.

| Window | Time | Observed GD | Mean GD from the permutation distribution | 95% CI for GD from the permutation distribution | Probability of the Observed GD statistic (or greater) under the permutation distribution |
| --- | --- | --- | --- | --- | --- |
| W1 | 100ms | 3.22 | 2.46 | 1.92, 3.14 | 0.04 |
| W2 | 156-160ms | 2.5 | 1.96 | 1.58, 2.43 | 0.04 |
| W3 | 380-400ms | 2.4 | 1.89 | 1.55, 2.3 | 0.02 |
| W4 | 424-484ms | 2.33 | 1.8 | 1.48, 2.19 | 0.03 |
| W5 | 580-604ms | 2.2 | 1.81 | 1.51, 2.17 | 0.04 |
| W6 | 660-800ms | 2.66 | 2.05 | 1.72, 2.4 | 0.01 |

In order to identify significant electrode clusters driving the topographic differences during these periods, a spatio-temporal permutation-based analysis was performed in BESA Statistics 2.0 for each of these time windows. An examination of the first of these periods to pass the window threshold test (window 4) revealed three significant clusters of electrodes that showed significantly different activity from Pretest to Posttest. The first significant cluster (*p* < 0.0001) was centered between the Cz and Fz electrodes and comprised the following EGI electrodes (approximate 10-10 equivalent is included if available (Luu & Ferree, 2005)): 5, 6 (Fcz), 7, 20, 30, 55 (CpZ), 106, 112 (FC2), 118. The second significant cluster (*p* = 0.0214) in window 4 was located over the FPZ region and included the following EGI electrodes (approximate 10-10 equivalent is included if available): 14 (FPZ), 15 (Fcz), 16 (Afz), 17, 21, 22 (FP1)). The final significant cluster (*p* = 0.022) in window 4 was found over the frontal-temporal region and comprised the following EGI electrodes (approximate 10-10 equivalent is included if available): 39, 40 (See Figure 4, panel B). The analysis of the second time period to pass the window threshold test (window 6), revealed one significant cluster of electrodes (*p* = 0.04) that centered near FT9 and F9 electrodes. This cluster consisted of EGI electrodes (approximate 10-10 equivalent is included if available): 48, 128 and 127 (See Figure 4, panel W1, W2, W3, W4, W5 and W6).

#### 3.2.3a Analysis of source difference for Generalized Learning

In order to estimate the source activations that lead to the map difference in the generalized learning condition that spanned from 252ms to 272ms, average distributed source images obtained from LAURA modeling were contrasted between Pretest and Posttest using a cluster-based permutation test in BESA Statistics 2.0 (Contrast: Posttest-Pretest). For each identified source cluster, the closest cortical region was determined using the MATLAB toolbox version of the Brede Database (Nielsen, 2003). For generalized learning, four clusters were identified as decreasing in activity following training: a cluster near superior temporal gyrus (31.5, - 30.9, 9.7, p = 0.231), a cluster in left superior parietal (−17.5, −79.9, 37.7, p = 0.335), a cluster in left anterior cingulate gyrus (−17.5, 11.1, 23.7, p = 0.39) and a cluster in right anterior cingulate gyrus (10.5, 25.1, 9.7, p = 0.559). While these four clusters do not pass the significance threshold of p<0.05, these identified regions directly align with areas that would be expected given our hypothesis that generalized learning of a difficult-to-understand talker helps to alleviate attention by reorganizing attention to the most diagnostic phonological features for the talker.

#### 3.2.3b Analysis of source difference for Rote Learning

Similar to Generalized Learning, we estimated the source activations that led to the six identified map differences (W1, W2, W3, W4, W5, and W6) in the rote learning condition by contrasting Pretest and Posttest average distributed source images obtained from LAURA modeling using a cluster-based permutation test in BESA Statistics 2.0 (Contrast: Posttest-Pretest). For each identified source cluster, the closest cortical region in Talairach coordinate space was determined using the MATLAB toolbox version of the Brede Database (Nielsen, 2003). Table 2 reports the results of the cluster-based permutation analysis for each of the six windows identified through the RAGU TANOVA analysis. It is important to note that the areas implicated by this analysis in the rote learning condition are to some extent consistent with episodic learning models (e.g. Spaniol et al., 2009).

**Table 2.** Results of the cluster-based permutation analysis on the average distributed source images obtained from LAURA modeling comparing Pretest and Posttest for each of the six windows identified through the RAGU TANOVA analysis for the Rote Learning Condition. For each identified cluster, the x,y,z location of peak activation is reported in Talairach coordinate space. Shaded rows indicate windows that pass the duration threshold in RAGU TANOVA analysis.

| Window | Time | Sources | Lobar anatomy | Cluster significance | Pretest activity | Posttest activity |
| --- | --- | --- | --- | --- | --- | --- |
| W1 | 100ms | -17.5, -51.9, -25.3 | Left cerebellum | 0.01 | 0.137 | 0.1026 |
|  |  | 31.5, 60.1, 23.7 | Right middle frontal gyrus | 0.02 | 0.173 | 0.1302 |
|  |  | 59.5, -51.9, -18.3 | Right inferior temporal gyrus | 0.07 | 0.1237 | 0.9824 |
|  |  | -31.5, 4.1, 16.7 | Left inferior frontal gyrus | 0.11 | 0.1821 | 0.1469 |
| W2 | 145-160ms | 24.5, -23.9, 16.7 | Right superior temporal gyrus | <.0001 | 0.1855 | 0.1406 |
|  |  | 10.5, -79.9, 44.7 | Right medial parietal/precuneus | 0.44 | 0.0327 | 0.0263 |
| W3 | 380-400ms | 10.5, -30.9, 9.7 | Right posterior cingulate | 0.18 | 0.2578 | 0.185 |
|  |  | 10.5, 32.1, 16.7 | Right anterior cingulate | 0.31 | 0.5819 | 0.4024 |
|  |  | 52.5, 18.1, 9.7 | Right inferior frontal gyrus | 0.48 | 0.1492 | 0.1135 |
| W4 | 424-484ms | 24.4, -23.9, 16.7 | Right superior temporal gyrus | 0.14 | 0.2937 | 0.2148 |
|  |  | -31.5, -9.9, 9.7 | Left temporal insula | 0.38 | 0.3886 | 0.3078 |
| W5 | 580-604ms | -31.5, -23.9, 9.7 | Left Heschl's gyrus | 0.07 | 0.4862 | 0.3298 |
|  |  | -17.5, -72.9, 30.7 | Left medial parietal/precuneus | 0.2 | 0.212 | 0.1597 |
| W6 | 660-800ms | -31.5, -2.9, 23.7 | Left inferior frontal gyrus | 0.23 | 0.4096 | 0.3171 |
|  |  | -3.5, -79.9, -4.3 | Lingual gyrus | 0.29 | 0.1262 | 0.1626 |
|  |  | -24.4, -65.9, 30.7 | Left precuneus | 0.31 | 0.2423 | 0.1797 |

## 4. Discussion

One model of generalization and abstraction in memory is that individual experiences are encoded as rote representations and that generalization emerges from the aggregate response of long-term memory given a novel test item. The test item elicits responses from prior experiences that are stored and the emergent response to the novel item is a generalization over those individual traces (e.g., McClelland and Rumelhart, 1985). If this were the case, there should be substantial similarities in rote learning and generalization learning with the primary difference being the strength of representation in rote learning (more instances of encoding the same trace). However, prior research has argued there are different mechanisms underlying rote and generalized perceptual learning of synthetic speech (e.g., Fenn et al., 2013) by showing different patterns of consolidation during sleep. The present patterns of neural responding in rote and generalized learning support and elaborate this view of learning. Generalization learning that goes beyond stimulus-specific experiences is marked by an amplitude reduction in the latter portion of the N1 wave into the peak of the P2 wave from 116ms to 208ms not seen in rote learning. These generalization training effects were followed later in the time-course of recognition by (1) a source configuration change from 250ms to 272ms that was estimated to arise from a decrease in activity in the right superior temporal gyrus, left superior parietal and the anterior cingulate gyrus bilaterally and (2) late negativity in the auditory evoked potential 580-800ms post-stimulus onset. Taken together, these changes suggest how generalized training is related to patterns of neural activity not seen in rote learning. Thus, this provides new information on how listeners transfer learning beyond stimulus-specific experiences and as such, offer one potential method listeners may use when adjusting to a novel talker. Unlike generalized learning, rote learning was only marked by a series of source configuration changes mostly occurring 380ms after stimuli onset. The demonstration that we observed changes in the N1-P2 complex for generalized learning and not for rote learning supports our view that the transfer of learning beyond talker specific experiences is garnered through an attentional reorganization process that adaptively reorganizes early auditory processing to cope with systematic acoustic variability at the level of a talker.

For example, the amplitude reduction we observed could reflect a sparsening of the memory code or stimulus representation for the synthetic talker’s speech. On the one hand, this could have decreased the redundancy of neurons responding to different stimuli. The decrease of auditory neural responses following training could account for a change in attentional processing such as that reported by Francis et al. (2000) showing that feedback during training shifts attention away from some acoustic cues and to others. Given that prior to learning, listeners may have been attending to relevant and irrelevant cues, the sharpening of the auditory neural representation of the learned speech would necessary shift attention to those properties that remain encoded neurally. It is also possible that the reduction in neural response reflects a kind of automatization (see Shiffrin and Schneider, 1977) of sensory processing. Training on this kind of synthetic speech does reduce working memory demands consistent with increased automatization and reduced cognitive load (Francis and Nusbaum, 2009). However, one might expect that if automatization were the explanation of the N1-P2 change observed in generalization learning, the same change should have been observed (which it was not) for rote learning given that this is much more consistent with the conditions for automatization whereas the diversity of stimuli presented in generalization learning is the antithesis of the conditions for automatization. Regardless, the present study cannot distinguish which of the mechanisms outlined are responsible for the changes in attention suggested by the N1-P2 complex observed for generalized learning. However whether attention is shifted to change the sensory encoding of stimuli, whether there is a reduction in the neural representation of learned stimuli due to the focus on relevant acoustic-phonetic cues, or whether the encoding of stimuli has become automatized, generalized learning results in changes in attentional processing heretofore not clearly linked to neural processes.

As previously discussed, the auditory evoked N1 potential has been argued to be composed of at least two physically, and arguably functionally distinct sources (Jääskeläinen et al., 2004; McEvoy et al., 1997; Picton, 2011), with the temporally earlier N1 source supporting mechanisms by which novel, unattended sounds are brought into awareness, while the temporally later N1 source supports additional attentional focus to features comprising the auditory object (Jääskeläinen et al., 2004). Our GFP analysis in the generalized learning condition revealed a significant change in the later part of the N1 time period, from 116ms to 208ms, starting approximately at the height of the N1 peak and lasting through to the P2 component. The absence of change in the temporally earlier N1 source and presence of the change during the temporally later N1 source is consistent with the view that generalized learning of synthetic speech is related to a substantial reduction in the demands of attention toward features comprising an auditory object (Gutschalk et al., 2008a; Jääskeläinen et al., 2004; Tiitinen et al., 1994). This interpretation is also consistent with cognitive models of generalized perceptual learning in the context of speech recognition, as generalized learning of synthetic speech has been described as a reduction of processing effort due to a more selective allocation of attention to phonetically informative features following training (Francis et al., 2000; Francis and Nusbaum, 2002; Schwab et al., 1985). The absence of a similar decrease in N1 following rote training supports the idea that rote learning in this setting does not substantially alter early attentional processes and as such may be much more similar to memory encoding of episodic traces. Consequently, it also offers an explanation as to why transfer of learning is found to a much greater extent following generalized training compared to rote training in the context of learning a difficult-to-understand talker.

Changes found in the P2 component of the auditory evoked potential in the generalized learning condition differ from those found in previous studies examining perceptual learning (Ross and Tremblay, 2009; Tremblay et al., 2014). While previous studies demonstrate post-sleep increases in the P2 component following training, generalized learning here, coincided with an immediate reduction in the P2 component in terms of its amplitude (see 3.2.2a). Given the reliance on sleep to consolidate rapidly acquired learning into long-term representations (c.f. Nusbaum et al., 2018) previous studies have argued that the sleep-dependent P2 change marks the consolidation of a feature-based representation in long-term memory (Ross and Tremblay, 2009; Tremblay et al., 2014). Here, we point out an implication of this interpretation: if the P2 component is sensitive to the formation of an additional featural representation in long-term memory, it indicates that the auditory P2 evoked response may be sensitive to the number of active featural representations serving current recognition. Under this view, the change observed in the auditory P2 evoked response following generalized training and not for rote learning, indicates that generalized training decreases the number of active featural representations required for on-going perception. Beyond the observed change in GFP following generalized training during the first half of the P2 response due to generalized training, the TANOVA analysis indicated a change in topography in the latter half of the P2 component between 250ms to 272ms (see 3.2.3a). While this window did not pass the duration threshold for RAGU, its appearance during the N1-P2 complex arguably elevates its relevance, and as such we interpret its appearance. The distributed source modeling with LAURA estimates that this window of topographic change was driven by a decrease in activity in the right superior temporal gyrus, left superior parietal and the anterior cingulate gyrus bilaterally. Both superior parietal cortex as well as the anterior cingulate have been argued to be responsible for attentional resource allocation for ongoing processing (Myers and Theodore, 2017; Piai et al., 2013; Wong et al., 2004). Further, the superior temporal gyrus has been associated with processing that is sensitive to talker-specific phonology (Myers and Theodore, 2017; Wong et al., 2004). These regions therefore appear to comprise a network capable of reorganizing attentional resources to acoustic cues, which are most meaningful for a to-be-learned talker. Taken collectively, the observed changes during the P2 window in the current study provide strong evidence that the transfer of learning beyond utterance-specific experiences is accomplished by reorganizing attention towards features that are most informative. Again, the lack of similar changes in the P2 component following rote training supports the idea that rote learning (at least in the context of understanding a difficult-to-understand talker) does not substantially alter early attentional processes and as such may be best thought of as a process that involves the simple formation of memory representations or associations between phonetic patterns and their meanings. Results from distributed source modeling with LAURA at the windows of topographic changes found in the rote learning condition appear to support this view, with many of the identified areas (see Table 2) implicated in episodic learning models (e.g. Spaniol et al., 2009).

Beyond changes in the N1-P2 complex, we observed a difference in the late negativity in the auditory evoked potential starting 580ms post-stimulus onset following generalized learning but not following rote learning. This mirrors results by Tremblay, Ross, Inoue, McClannahan, and Collet (2014), who found late negativity in the auditory evoked potential starting 600ms post-stimulus onset following training. As previously mentioned, this late potential may reflect improvements in trial-by-trial, prediction-error monitoring that drives the reorganization or formation of perceptual categories (Ashby et al., 1998; Ashby and O’Brien, 2005). Indeed, trial-by-trial prediction and error correction processes are thought to play a critical part in implicit category learning, as such processes are thought to provide the feedback critical to direct learning. This post-training modulation of the late negative response was only found following generalized learning and not rote learning which may serve generalized training for better transfer of learning compared to rote training.

## 5. Conclusions

Previous research has suggested that generalization and rote learning may be mediated by different neural mechanisms. The present study tested this directly by comparing how patterns of neural responses during speech recognition change following rote and generalization learning. The present results demonstrate substantial differences in neural responses for these two types of learning. On the one hand, rote and generalized learning might have been supported by the same neural process of encoding a series of auditory traces corresponding to experienced spoken words and then recognition, whether of a previously heard word or a new word, would either retrieve the trace of the learned word or an aggregate response over similar words thereby generalizing on the fly. While this corresponds to an entire class of memory models, the present data reject this view showing that generalization learning entails early neurosensory changes in processing that may be attributed to changes in attention that are not seen in rote learning. This difference argues against a passive, bottom-up fixed speech processing system that simply records auditory experiences and supports the view that speech perception is mediated by active neural processing, by which listeners can learn to selectively attend to the acoustic-phonetic characteristics of a novel talker. Moreover, this view of speech perception that depends on the active use of general attentional mechanisms to adaptively respond to the challenge of acoustic-phonetic uncertainty may be a very general system used to deal with the problem of pattern variability and recognition to maintain robust phonetic constancy when noise, distortion, or other acoustic challenges occur.

## Acknowledgments

The authors thank Sophia Uddin for helpful comments on an earlier draft, as well as Nina Bartram, Edward Wagner, and Brendan Colson for assistance with data collection.

## References

Ahveninen, J., Jääskeläinen, I.P., Raij, T., Bonmassar, G., Devore, S., Hämäläinen, M., Levänen, S., Lin, F.H., Sams, M., Shinn-Cunningham, B.G., Witzel, T., Belliveau, J.W., 2006. Task-modulated “what” and “where” pathways in human auditory cortex. Proc. Natl. Acad. Sci. U. S. A. 103, 14608–14613. https://doi.org/10.1073/pnas.0510480103

Alain, C., Campeanu, S., Tremblay, K., 2010. Changes in sensory evoked responses coincide with rapid improvement in speech identification performance. J. Cogn. Neurosci. 22, 392–403. https://doi.org/10.1162/jocn.2009.21279

Alain, C., Snyder, J.S., 2008. Age-related differences in auditory evoked responses during rapid perceptual learning. Clin. Neurophysiol. 119, 356–366. https://doi.org/10.1016/j.clinph.2007.10.024

Alain, C., Snyder, J.S., He, Y., Reinke, K.S., 2007. Changes in auditory cortex parallel rapid perceptual learning. Cereb. Cortex 17, 1074–1084. https://doi.org/10.1093/cercor/bhl018

Andrews, R.J., Knight, R.T., Kirby, R.P., 1990. Evoked potential mapping of auditory and somatosensory cortices in the miniature swine. Neurosci. Lett. 114, 27–31. https://doi.org/10.1016/0304-3940(90)90423-7

Ashby, F.G., Alfonso-Reese, L. a, Turken, a U., Waldron, E.M., 1998. A neuropsychological theory of multiple systems in category learning. Psychol. Rev. 105, 442–81. https://doi.org/10.1037/0033-295X.105.3.442

Ashby, F.G., O’Brien, J.B., 2005. Category learning and multiple memory systems. Trends Cogn. Sci. https://doi.org/10.1016/j.tics.2004.12.003

Berg, P., Scherg, M., 1994. A multiple source approach to the correction of eye artifacts. Electroencephalogr. Clin. Neurophysiol. 90, 229–241. https://doi.org/10.1016/0013-4694(94)90094-9

Bosnyak, D.J., Eaton, R.A., Roberts, L.E., 2004. Distributed auditory cortical representations are modified when non-musicians are trained at pitch discrimination with 40 Hz amplitude modulated tones. Cereb. Cortex 14, 1088–1099. https://doi.org/10.1093/cercor/bhh068

Chomsky, N., Halle, M., 1968. The Sound Pattern of English., The MIT Press.

Egan, J.P., 1948. Articulation testing methods. Laryngoscope 58, 955–991. https://doi.org/10.1288/00005537-194809000-00002

Fenn, K.M., Margoliash, D., Nusbaum, H.C., 2013. Sleep restores loss of generalized but not rote learning of synthetic speech. Cognition 128, 280–286. https://doi.org/10.1016/j.cognition.2013.04.007

Fenn, K.M., Nusbaum, H.C., Margoliash, D., 2003. Consolidation during sleep of perceptual learning of spoken language. Nature. https://doi.org/10.1038/nature01951

Francis, A.L., Baldwin, K., Nusbaum, H.C., 2000. Effects of training on attention to acoustic cues. Percept. Psychophys. 62, 1668–1680. https://doi.org/10.3758/BF03212164

Francis, A.L., Nusbaum, H.C., 2009. Effects of intelligibility on working memory demand for speech perception. Attention, Perception, Psychophys. 71, 1360–1374. https://doi.org/10.3758/APP.71.6.1360

Francis, A.L., Nusbaum, H.C., 2002. Selective attention and the acquisition of new phonetic categories. J. Exp. Psychol. Hum. Percept. Perform. 28, 349–366. https://doi.org/10.1037//0096-1523.28.2.349

Goldstone, R.L., 1998. Perceptual learning. Annu. Rev. Psychol. 49, 585–612. https://doi.org/10.1146/annurev.psych.49.1.585

Greenspan, S.L., Nusbaum, H.C., Pisoni, D.B., 1988. Perceptual learning of synthetic speech produced by rule. J. Exp. Psychol. Learn. Mem. Cogn. 14, 421–433. https://doi.org/10.1037/0278-7393.14.3.421

Gutschalk, A., Micheyl, C., Oxenham, A.J., 2008a. Neural Correlates of Auditory Perceptual Awareness under Informational Masking. PLoS Biol. 6, e138. https://doi.org/10.1371/journal.pbio.0060138

Gutschalk, A., Micheyl, C., Oxenham, A.J., Kriegstein, K. von, Warren, J., 2008b. Neural Correlates of Auditory Perceptual Awareness under Informational Masking. PLoS Biol. 6, e138. https://doi.org/10.1371/journal.pbio.0060138

Heald, S.L.M., Nusbaum, H.C., 2014. Speech perception as an active cognitive process. Front. Syst. Neurosci. 8, 35. https://doi.org/10.3389/fnsys.2014.00035

Ing-Simmons, N., 1994. RSYNTH: Complete speech synthesis system for UNIX.

Jääskeläinen, I.P., Ahveninen, J., Bonmassar, G., Dale, A.M., Ilmoniemi, R.J., Levanen, S., Lin, F.-H., May, P., Melcher, J., Stufflebeam, S., Tiitinen, H., Belliveau, J.W., 2004. Human posterior auditory cortex gates novel sounds to consciousness. Proc. Natl. Acad. Sci. 101, 6809–6814. https://doi.org/10.1073/pnas.0303760101

Klatt, D.H., 1980. Software for a cascade/paralell formant synthesizer. J. Acoust. Soc. Am. 67, 971–995. https://doi.org/10.3758/BRM.42.3.863

Koenig, T., Kottlow, M., Stein, M., Melie-García, L., 2011. Ragu: A free tool for the analysis of EEG and MEG event-related scalp field data using global randomization statistics. Comput. Intell. Neurosci. 2011, 1–14. https://doi.org/10.1155/2011/938925

Liberman, A.M., 1970. The grammars of speech and language. Cogn. Psychol. 1, 301–323. https://doi.org/10.1016/0010-0285(70)90018-6

Liégeois-Chauvel, C., Musolino, A., Badier, J.M., Marquis, P., Chauvel, P., 1994. Evoked potentials recorded from the auditory cortex in man: evaluation and topography of the middle latency components. Electroencephalogr. Clin. Neurophysiol. Evoked Potentials 92, 204–214. https://doi.org/10.1016/0168-5597(94)90064-7

Luu, P., Ferree, T.C., 2000. Determination of the Geodesic Sensor Nets’ Average Electrode Positions and Their 10-10 International Equivalents Sleep actigraphy View project C elegans View project.

McCallum, W.C., Curry, S.H., 1979. Hemisphere Differences in Event Related Potentials and CNV’s Associated with Monaural Stimuli and Lateralized Motor Responses, in: Human Evoked Potentials. Springer US, pp. 235–250. https://doi.org/10.1007/978-1-4684-3483-5_16

McClelland, J.L., Rumelhart, D.E., 1985. Distributed Memory and the Representation of General and Specific Information. J. Exp. Psychol. Gen. 114, 159–188. https://doi.org/10.1037/0096-3445.114.2.159

McEvoy, L., Levänen, S., Loveless, N., 1997. Temporal characteristics of auditory sensory memory: Neuromagnetic evidence. Psychophysiology 34, 308–316. https://doi.org/10.1111/j.1469-8986.1997.tb02401.x

Murray, M.M., Brunet, D., Michel, C.M., 2008. Topographic ERP analyses: A step-by-step tutorial review. Brain Topogr. 20, 249–264. https://doi.org/10.1007/s10548-008-0054-5

Myers, E.B., Theodore, R.M., 2017. Voice-sensitive brain networks encode talker-specific phonetic detail. Brain Lang. 165, 33–44. https://doi.org/10.1016/j.bandl.2016.11.001

Näätänen, R., Winkler, I., 1999. The concept of auditory stimulus representation in cognitive neuroscience. Psychol. Bull. 125, 826–859. https://doi.org/10.1037/0033-2909.125.6.826

Nielsen, F.Å., 2003. The Brede database: a small database for functional neuroimaging. Neuroimage 19, 19–22.

Nosofsky, R.M., 1986. Attention, Similarity, and the Identification-Categorization Relationship. J. Exp. Psychol. Gen. 115, 39–57. https://doi.org/10.1037/0096-3445.115.1.39

Nusbaum, H.C., Pisoni, D.B., 1985. Constraints on the perception of synthetic speech generated by rule, Behavior Research Methods, Instruments, & Computers.

Nusbaum, H.C., Uddin, S., Van Hedger, S.C., Heald, S.L., 2018. Consolidating skill learning through sleep. Curr. Opin. Behav. Sci. 20. https://doi.org/10.1016/j.cobeha.2018.01.013

Petkov, C.I., Kang, X., Alho, K., Bertrand, O., Yund, E.W., Woods, D.L., 2004. Attentional modulation of human auditory cortex. Nat. Neurosci. 7, 658–663. https://doi.org/10.1038/nn1256

Piai, V., Roelofs, A., Acheson, D.J., Takashima, A., 2013. Attention for speaking: domain-general control from the anterior cingulate cortex in spoken word production. Front. Hum. Neurosci. 7, 832. https://doi.org/10.3389/fnhum.2013.00832

Picton, T.W., 2011. Human auditory evoked potentials, Ear and Hearing. Plural Publishing, San Diego.

Picton, T.W., Van Roon, P., Armilio, M.L., Berg, P., Ille, N., Scherg, M., 2000. The correction of ocular artifacts: A topographic perspective. Clin. Neurophysiol. 111, 53–65. https://doi.org/10.1016/S1388-2457(99)00227-8

Rauschecker, J.P., Tian, B., 2000. Mechanisms and streams for processing of “what” and “where” in auditory cortex. Proc. Natl. Acad. Sci. 97, 11800–11806. https://doi.org/10.1073/pnas.97.22.11800

Reinke, K.S., He, Y., Wang, C., Alain, C., 2003. Perceptual learning modulates sensory evoked response during vowel segregation. Cogn. Brain Res. 17, 781–791. https://doi.org/10.1016/S0926-6410(03)00202-7

Ross, B., Jamali, S., Tremblay, K.L., Abdi, H., Toga, A., Evans, A., Preissl, H., Leahy, R., Sams, M., Shinn-Cunningham, B., 2013. Plasticity in neuromagnetic cortical responses suggests enhanced auditory object representation. BMC Neurosci. 14, 151. https://doi.org/10.1186/1471-2202-14-151

Ross, B., Tremblay, K., 2009. Stimulus experience modifies auditory neuromagnetic responses in young and older listeners. Hear. Res. 248, 48–59. https://doi.org/10.1016/j.heares.2008.11.012

Russell, G.S., Jeffrey Eriksen, K., Poolman, P., Luu, P., Tucker, D.M., 2005. Geodesic photogrammetry for localizing sensor positions in dense-array EEG. Clin. Neurophysiol. 116, 1130–1140. https://doi.org/10.1016/j.clinph.2004.12.022

Scherg, M., Vajsar, J., Picton, T.W., 1989. A Source Analysis of the Late Human Auditory Evoked Potentials. J. Cogn. Neurosci. 1, 336–355. https://doi.org/10.1162/jocn.1989.1.4.336

Schwab, E.C., Nusbaum, H.C., Pisoni, D.B., 1985. Some effects of training on the perception of synthetic speech. Hum. Factors 27, 395–408. https://doi.org/10.1177/001872088502700404

Shahin, A., Roberts, L.E., Pantev, C., Trainor, L.J., Ross, B., 2005. Modulation of P2 auditory-evoked responses by the spectral complexity of musical sounds. Neuroreport 16, 1781–1785. https://doi.org/10.1097/01.wnr.0000185017.29316.63

Shiffrin, R.M., Schneider, W., 1977. Controlled and automatic human information processing: II. Perceptual learning, automatic attending and a general theory. Psychol. Rev. 84, 127–190. https://doi.org/10.1037/0033-295X.84.2.127

Spaniol, J., Davidson, P.S.R., Kim, A.S.N., Han, H., Moscovitch, M., Grady, C.L., 2009. Event-related fMRI studies of episodic encoding and retrieval: Meta-analyses using activation likelihood estimation. Neuropsychologia. https://doi.org/10.1016/j.neuropsychologia.2009.02.028

Tiitinen, H., May, P., Reinikainen, K., Näätänen, R., 1994. Attentive novelty detection in humans is governed by pre-attentive sensory memory. Nature 372, 90–92. https://doi.org/10.1038/372090a0

Tremblay, K.L., Ross, B., Inoue, K., McClannahan, K., Collet, G., 2014. Is the auditory evoked P2 response a biomarker of learning? Front. Syst. Neurosci. 8, 28. https://doi.org/10.3389/fnsys.2014.00028

Tremblay, K.L., Shahin, A.J., Picton, T., Ross, B., 2009. Auditory training alters the physiological detection of stimulus-specific cues in humans. Clin. Neurophysiol. 120, 128–35. https://doi.org/10.1016/j.clinph.2008.10.005

Weatherholtz, K., Jaeger, T.F., 2016. Speech perception and generalization across speakers and accents. Linguist. Oxford Res. Encycl. https://doi.org/10.1093/acrefore/9780199384655.013.95

Wong, P.C.M., Nusbaum, H.C., Small, S.L., 2004. Neural bases of talker normalization. J. Cogn. Neurosci. 16, 1173–1184. https://doi.org/10.1162/0898929041920522

Woods, D.L., Stecker, G.C., Rinne, T., Herron, T.J., Cate, A.D., Yund, E.W., Liao, I., Kang, X., 2009. Functional maps of human auditory cortex: Effects of acoustic features and attention. PLoS One 4. https://doi.org/10.1371/journal.pone.0005183

Yvert, B., Fischer, C., Bertrand, O., Pernier, J., 2005. Localization of human supratemporal auditory areas from intracerebral auditory evoked potentials using distributed source models. Neuroimage 28, 140–153. https://doi.org/10.1016/j.neuroimage.2005.05.056

